# Sorting *F. graminearum* core effector candidates shows multiple fungal proteins that target the wheat cell nucleus during Fusarium Head Blight

**DOI:** 10.1101/2025.02.05.636710

**Authors:** Shimlal Ayilalath, Lilian Faurie, Emmanuel Vanrobays, Florian Rocher, Loriane Loizeau, Géraldine Philippe, Marie Javel, Mickaël Bosio, Christophe Sallaud, Christophe Tatout, Ludovic Bonhomme

## Abstract

Effectors are small molecules secreted by microbial pathogens that disrupt host basal functioning and responses during infection by targeting various plant susceptibility factors. This study reports a candidate selection approach for identifying novel, potential plant nuclear localized effectors from *Fusarium graminearum* secretory proteins. From a dataset of core secretory proteins conserved across several *Fusarium* strains, candidates were selected based on predicted nuclear localization, structural characteristics, and expression profiles during infection. Transient expression in *Nicotiana benthamiana* confirmed accumulation in the plant nucleus, that were further confirmed in wheat protoplasts. One of these proteins was selected for yeast two-hybrid (Y2H) screening to identify wheat protein targets, using a *Fusarium*-infected wheat spike cDNA library specifically generated for this study. The screening identified a high confident interaction with a nuclear-localized wheat beta-amylase 2. The structural modeling of the protein complex between beta-amylase 2 and the putative effector was used to predict interacting amino acid residues and informed a deletion analysis to disrupt the interaction. This research identifies a *F. graminearum* secretory core protein that interacts with a conserved wheat beta-amylase 2, showcasing a method to select pathogenicity factors conserved across multiple pathogens and host plants, with implications for developing broad-spectrum resistance strategies.

**Highlight:** This study proposes a method based on *in silico* and *in vivo* screens to identify interacting pairs between core pathogenicity effectors localized in the nucleus and susceptibility factors in plants.

## Introduction

Effectors are small molecules produced by microbial pathogens that interfere with a wide array of host cellular responses to promote conducive conditions for pathogen development and enhance infection processes. Secreted and then released into the host tissues, the effectors reach precise subcellular targets to achieve their roles through direct or indirect interactions with molecular functions and/or cellular components, the so-called plant susceptibility factors (Zhang *et al*., 2022). In plant-pathogen interactions, these targets are remarkably diverse, depicting the need of the pathogen to control multiple specialized host functions at specific stages of the infection process (Deslandes and Rivas, 2011). The nucleus is obviously a prime target for many effectors because of its central role in regulating plant cell functions and responses and a wide range of plant pathogens produce nuclear effectors, such as bacteria, viruses, nematodes, fungi and oomycetes (Harris *et al*., 2023). All these studies have demonstrated their remarkable actions in manipulating plant cell responses, especially through direct or indirect effects on the expression of a diverse set of host genes to manipulate plant immunity, to suppress the activation of genes responsible for plant defense, and even to interfere with generic and specific pathogen recognition signals (Jiang *et al*., 2007; Zhang *et al*., 2022; Cai *et al*., 2023). For instance, in *Nicotiana benthamiana* (*N. benthamiana*), the bacterial effector Pi23226 from *Phytophthora infestans* has been proven to accumulate in the nucleus and to interfere with the 25S rRNA biogenesis. This ultimately results in the disruption of protein translation and promotes cell death, thereby enabling the pathogen to circumvent the plant’s defensive mechanisms and facilitate its proliferation within the plant (Lee *et al*., 2023). Similarly, during the rice blast disease, the fungal effector AvrBs3, secreted by *Magnaporthe oryzae*, has been shown to inhibit the action of certain/specific transcription factors by directly binding to the host DNA, which results in the suppression of the defense response (Zhang *et al*., 2022). As such, increasing our knowledge on nuclear-targeted microbial effectors will lead to a better understanding of plant pathogen interactions and help to further discover the range of susceptibility factors that pathogens are able to manipulate (Todd *et al*., 2022). By screening their precise actions, this can leverage new avenues to unveil the host vulnerabilities that pathogens exploit during infection and to identify new targeted strategies for crop protection. Fusarium head blight (FHB) is one of the most devastating plant diseases in small grains such as wheat and barley (Dean *et al*., 2012). Primarily caused by the fungus *Fusarium graminearum* (*F. graminearum*), FHB not only reduces grain yield and quality but also contaminates grains with harmful mycotoxins leading to serious health risk to humans and animals (Ekwomadu *et al*., 2021; Xu *et al*., 2023). Over the past 30 years, genetic approaches have yielded more than 650 QTLs for FHB resistance but providing overall small improvements in the fields with high dependency on environmental conditions (Steiner *et al*., 2017). Searching for fungal effectors is opening up new opportunities to identify novel key determinants of plant disease, further exploring related wheat susceptibility factors to identify new sources of resistance (Fabre *et al*., 2020). To date, about a dozen *F. graminearum* effectors have been identified with diverse functions and plant subcellular targets (Voigt *et al*., 2005; Jiang *et al*., 2020; Hao *et al*., 2023; Jin *et al*., 2024). For instance, OSP24 interacts with the *Triticum aestivum* Sucrose non-fermenting-1-related protein Kinase 1α (TaSnRK1α) to promote the degradation of the kinase by the proteasome, resulting in a decreased FHB resistance in wheat (Jiang *et al*., 2020). TaSnRK1α modulates the activity of the enzyme AGPase (ADP-glucose pyrophosphorylase), which is a key enzyme in starch biosynthesis linking essential metabolic functions such as sugar metabolism and resistance to fungal pathogens. Another effector, called FgNls1, contains several Nuclear Localization Signals (NLS) and its transient expression in *N. benthamiana* as a FgNls1-GFP fusion protein shows its accumulation in the nuclei of plant cells (Hao et al., 2023). Consistent with its nuclear localization, affinity chromatography experiments identified a wheat histone H2B as a main target protein that interacts with FgNls1. These few examples demonstrate the unequivocal role of *F. graminearum* effectors in wheat susceptibility to FHB and suggest that the nucleus can be a primary target to control an intricate set of plant responses. As such, the identification of interacting pairs between effectors and susceptibility factors is a promising strategy to weaken the FHB disease and to improve wheat tolerance.

In our previous works, we identified a core secretome of 357 proteins found to be expressed by different strains of *F. graminearum* of varying aggressiveness in different wheat cultivars of contrasting susceptibility (Rocher *et al*., 2022). Displaying close expression patterns that depicted two early waves of secretory protein delivery over a 96-hour period, these data showed that the genetic program developed by *F. graminearum* during FHB is well conserved across different strains and whatever the host’s genetic background. One set of secretory proteins, called early secretory proteins, is typically expressed during the very early stages of infection (i.e. until 48 hours after fungal inoculation) while late secretory proteins are expressed at later stages of infection (i.e. starting from 72 hours after inoculation) (Rocher *et al*., 2022). The conservation of these expression dynamics in different *F. graminearum* strains suggests that these secretory proteins play an active role in promoting infection by targeting essential host functions at specific times of the infection. Interestingly, among these core secretory proteins, 46 harbored plant NLS and 41 were found to be co-regulated with regulatory hubs of the complex gene regulation network of FHB responses in wheat (Rocher *et al*., 2024), suggesting their potential role in interfering with massive gene expression regulation. Based on this dataset, we aim to demonstrate that multiple *F. graminearum* secretory proteins effectively accumulate in the plant nucleus to fulfill their function. First, starting from the 357 core secretory proteins identified in Rocher *et al*. (2022), we selected a set of 14 candidates based on their structural features predicted by *in silico* tools and their early and late expression profiles during infection. Nuclear accumulation was assayed using transient expression in *N. benthamiana* and subsequently confirmed in wheat protoplasts for a subset of them. Bringing all this information together allowed the selection of one specific secreted protein hereafter called Effector18 (EFF18) to determine whether it could physically interact with wheat proteins, using the yeast two-hybrid (Y2H) system. For that we designed a specific wheat spike cDNA library and identified four targets with high confidence scores. While the identified interacting host proteins were further validated by yeast two hybrid interaction assay, we focused our attention to EFF18 and LOC123105923, a beta-amylase 2 protein, because their interaction was observed both in yeast and when using the *in silico* prediction tool ColabFold_Multimer. Finally, we took advantage of the predicted interacting complex to define the putative interacting residues and used these predictions to design a deletion analysis to confirm the predicted interaction domain. This study identifies a core secretory protein from *F. graminearum* that interacts with a well-conserved beta-amylase 2 and illustrates a strategy to select pathogenicity factors conserved across multiple pathogens and their host plants to develop promising generalized broad-spectrum resistance.

## Materials and Methods

### Microbial strains, plant materials, growth conditions and plant assays

The *F. graminearum* strain MDC_Fg1 considered aggressive on different European cultivars (Alouane *et al*., 2018; Fabre *et al*., 2021; Rocher *et al*., 2022) was used in this study. Spores were obtained as previously detailed (Rocher *et al*., 2022), kept frozen until use and adjusted to a concentration of 1×10^5^ spores/ml in distilled water before inoculation. The ‘Recital’ wheat cultivar, an FHB-susceptible winter wheat (Rocher *et al*., 2022, 2024), was used for transient expression assay and to prepare a specific cDNA library. For wheat protoplast preparation, surface sterilized seeds of the ‘Recital’ wheat cultivar were germinated on a filter paper in petri dish in a germination chamber at 25°C with a 16:8 light cycle for 3-5 days. Seedlings were then transferred in pots containing sterilized soil in a growth chamber with same growth conditions as above, for 7 days until plants reached the two-leaf stage. *Agrobacterium* infiltration assay was performed using *N. benthamiana* grown on sterilized soil and incubated in a germination chamber maintained at 25°C with a 16:8 light cycle. Seedlings were transferred to a growth chamber with the same conditions for 3-4 weeks until plants reached the 5-7 leaves stage appropriate for infiltration.

### Transient expression of *F. graminearum* secretory proteins in plant cells

Genomic DNA sequences from *F. graminearum* MDC_Fg1 encoding the 14 secretory proteins excluding their signal peptide sequences were used for *de novo* gene synthesis (Genscript USA Inc). The synthesized sequences were cloned into the pGWB605 destination vector (Nakagawa *et al*., 2007) to express the proteins as C-terminal GFP fusions under the control of the p35S promoter. A second pGWB605 destination vector was designed to express a NLS fused to the mRFP under the control of the pDon promoter. In addition, a third vector expressing the P19 suppressor of RNA silencing was used in all transient expression assay to increase the expression level of co-expressed proteins (Petersen *et al*., 2019). Transient expression assays were performed by *Agrobacterium* infiltration of *N. benthamiana* leaves and observed after 72 to 96 hours after infiltration as described in Sparkes *et al*. (2006). Localization of proteins was analyzed using a Zeiss Cell Observer Spinning Disk microscope under 40X objective. A second transient expression assay was performed in wheat protoplast using the Recital wheat cultivar and the polyethylene glycol-mediated method as previously described (Luo *et al*., 2022). Protoplast viability was assessed with fluorescent visibility stain (EVANS Blue) and observed under a wide field microscope under 40X objective and 50,000 protoplasts were mixed with the secretory protein-GFP fusion plasmid and NLS-mRFP. The transfected protoplasts were incubated at 25°C for 2 days and observed using a Zeiss LSM 800 confocal microscope under a 63X oil immersion objective. The NLS-mRFP nuclear marker was used to determine the nuclear localization of the secretory proteins.

### Yeast 2-Hybrid assays

Protein-protein interactions were performed by Hybrigenics services (Evry, France) using the ULTImate Yeast 2-hybrid (Y2H) system and a dedicated cDNA library obtained from Recital wheat cultivar inoculated with the *F. graminearum* strain MDC_Fg1. To prepare the cDNA library, FHB-infected and mock spikes at different stages of the infection dynamics were produced as previously detailed (Rocher *et al*., 2022). Briefly, plants were sawn in buckets, vernalized during six weeks and transferred in a growth chamber with controlled conditions (16:8 light cycle with 21°C/17°C (day/night) temperatures and relative humidity of 80%). Infection was performed at mid-anthesis by inoculating the floral cavity of three synchronous flowering spikes. Spikelets inoculated with water served as controls. For each individual, the inoculated spikelets were collected and immediately frozen in liquid nitrogen. The cDNA library was constructed using inoculated spikelets collected from four plants at 48 and 72hpi as in Sasi et al. (Sasi *et al*., 2023). The resulting library was cloned into the Y2H vector as a pray construct. Coding sequencing of the secretory proteins without the signal peptide were cloned into Y2H vectors as bait constructs and expressed as a fusion protein with the LexA and/ or Gal4 DNA-binding domain. The bait constructs were transformed into yeast strain AH109 and the Y2H screening was performed by mating with the yeast strain Y187 containing the prey, which corresponds to the wheat cDNA fused to the DNA activation domain. Interaction-dependent activation of the reporter genes allowed for the identification of positive interactions. Identified binding partners in Y2H screens were classified in 4 grades (A, B, C and D) using the Hybrigenics Possible Biological Score (PBS) scoring system reflecting the reliability of the interactions where grade A represents the strongest interactions that exhibit high specificity, and which have been observed in independent experiments (Velásquez-Zapata *et al*., 2021). Selected interactions of grade A and B were subsequently validated using the Yeast 2-hybrid from Clontech, MATCHMAKER GAL4 Two-Hybrid System. Entry vectors containing bait and prey were synthesized by Genscript USA Inc and subsequently subcloned into the bait vector pDEST-GBKT7 or the pray vector pDEST-GADT7 (Rossignol *et al*., 2007). Yeast cultures were grown at 30 °C on Yeast Extract Peptone Dextrose (YPD) or on selective Synthetic Defined (SD) media. *Saccharomyces cerevisiae* haploid strains AH109 Gold and Y187 were transformed according to the classical protocol (Gietz and Woods, 2002), respectively with the secretory proteins cloned into the bait vector pDEST-GBKT7 or into the effectors cloned into the prey vector pDEST-GADT7 and grown on appropriate medium (SD-Trp or SD-Leu plates). After mating on YPD and selection of diploids on SD-Leu-Trp medium, interactions were tested on stringent medium SD-Leu-Trp-His-Ade. Empty pDEST-GBKT7 or pDEST-GADT7 vectors were used as negative controls. Two deletion derivatives were designed for EFF18, a C-terminal deletion and deletion of 45 amino acids corresponding to interacting residues of EFF18 with LOC123105923. C-terminal deletion called EFF18_DC was carried out with Phusion PCR DNA Polymerase (Thermo Fischer Scientific) on pUC57_EFF18 with primers C-term_EFF18_Forward (5’- GGGGACAAGTTTGTACAAAAAAGCAGGCTTCATGGTGCCCAAGGCTGACA-3’) and C-term-EFF18_Reverse (5’- CGGGGACCACTTTGTACAAGAAAGCTGGGTGTTAGTAGCCGAAAGGCGAAAGG- 3’). Deletion of interacting residues (EFF18_DM) was carried out on pUC57_EFF18 by using QuikChange II Site-Directed Mutagenesis Kit (Agilent) with primers Interaction_EFF18_Forward (5’- AATACATCCTTTCGCCTTTCGGCTAC<u>ACCGGCGCGGCGCCTGACCA</u>-3’) and Interaction_EFF18_Reverse (5’- <u>TGGTCAGGCGCCGCGCCGGTGTAGCCGAAAGGCGAAAGGATGTATT</u>-3’) designed by overlapping the deletion area (underlined). PCR products were subsequently introduced in pGADT7 and used as bait for Y2H assay.

### *In silico* analyses

Subcellular localization prediction of secreted proteins was carried out using the web tool LOCALIZER1.0.4 (Sperschneider *et al*., 2017) from mature protein sequences. SignalP5.0 was used to define the signal peptide in the secreted proteins (Nielsen *et al*., 2019). Protein features of sequences without the signal peptide were evaluated through the EffectorP-fungi 3.0 web tool (Sperschneider and Dodds, 2022). Depicter2 server was used to predict Intrinsically Disordered Region (IDR) (Basu *et al*., 2023). Monomeric and multimeric structural predictions of candidate proteins were computed with ColabFold Batch (v.1.5.5) (Mirdita *et al*., 2022). Configuration of ColabFold settings were maintained as default, with 5 predicted structural models, each comprising 3 cycles/reiterations per model. ColabFold scores calculated from monomeric or multimeric protein structures called pTM (Predicted Template Modeling) and ipTM (Interface Predicted Template Modeling) respectively, were used as global indicators of the prediction confidence. Alphafold-Multimer Local Interaction Score (AFM-LIS) (Kim *et al*., 2024) was used to predict protein interactions at the residual scale using the Local Interaction Score (LIS) and the Local Interaction Area (LIA). Probability of interaction of individual amino acids was based on the Predicted Aligned Error (PAE) scores calculated from the PAE matrix with a threshold of 12 Ångström (distance between amino acids). To predict positive protein protein interaction, the LIS and LIA were evaluated according to the following thresholds: a best LIS ≥ 0.203 and a best LIA ≥ 3432 (Kim *et al*., 2024). Visualization of 3D structures and identification of interacting residues at less than 5 Angstroms of distance were performed using ChimeraX (v.1.7.1) (Goddard *et al*., 2018). Clustering analyses were performed with FoldSeek (Van Kempen *et al*., 2024) using predicted protein structures as .pdb files available at RCSB Protein Data Bank (RCSB PDB) or predicted by ColabFold. FoldSeek was used with default parameters except for coverage set at 0.8, Predicted Local Distance Difference Test (pLDDT) threshold set at 0.7 and TM score threshold that quantifies the similarity between two protein structures set at 0.7. Clustering was performed either on the 351 core secreted proteins (pdf files were available for 351/357 accessions) alone or in combination with 3,991 additional putative effectors collected from the literature (Derbyshire and Raffaele, 2023; Seong and Krasileva, 2023; Rocafort *et al*., 2022). Genetic diversity was assessed using a set of 22 fungi species including basidiomycetes and ascomycetes and 22 plant species including unicellular algae, bryophytes and angiosperms. For each of the plant and fungi species, the protein database was retrieved from the NCBI/Taxonomy browser (**Supplemental Tables 1 and 2**). The search for orthologous sequences was carried out against downloaded databases of representative species of fungi (for secretory proteins) or plants (for wheat targets). For *F. graminearum* and wheat proteins, a tyrosine kinase protein (kinase, PF07714) (Alouane *et al*., 2021) and the Replication factor A protein 1 (RPA, YAR007C) (Cappela-Gutierrez *et al*., 2014) were selected as controls for their high and low percentage of identity, respectively, while AGAMOUS-LIKE 62 (AGL62, Q9FKK2) (Nam et al., 2004) and MUTS HOMOLOG 1 (MSH1, Q84LK0) (Bay and Guo, 2023) were chosen for similar reasons in wheat. The MMseqs2 tool (v.15.6f452) (Steinegger and Söding, 2017) was used with the reciprocal best hits (easy-rbh) option to select the best potential ortholog. Protein sequences were aligned using the mafft tool (v.7.525) (Katoh and Standley, 2013) and the output files were used as input for iqtree (v.2.3.3) by generating 1000 bootstraps. The treefile format was imported and modified using the iTOL web service (v.6) (Letunic and Bork, 2024). The Conserved Domain Database (CDD) web server (Lu *et al*., 2020) was used to identify conserved protein domains in protein sequences via Position-Specific Iterated BLAST (RPS-BLAST). The conserved domains present in the secreted proteins and their targets were identified using the CD-search batch option restricted to protein domains annotated in the pfam database (Mistry *et al*., 2021). Positions of conserved domains were used to draw the protein structures of secretory proteins and their putative targets identified by Y2H using the drawProtein R package. Bioinformatic scripts are available at https://gitlab.com/lilian_fau/fhb_secure/-/wikis/home.

## RESULTS

### *In silico* characterization of the core secretome of 357 proteins from *Fusarium graminearum*

As a starting point for this study, the 357 putative core secretory proteins previously identified by Rocher *et al*. (2022) (**Supplementary Table 3**) to further refine their characteristics and analyze their similarities both to one another and to those described in available published studies (Rocafort *et al*., 2022; Derbyshire and Raffaele, 2023; Seong and Krasileva, 2023). The 357 *F. graminearum* genes were systematically expressed and regulated during FHB infection in three strains of contrasting aggressiveness or when facing five different wheat host cultivars of contrasting susceptibility (Rocher *et al*., 2022). Overall, this gene set encodes small proteins, with an average size of 384 amino acids (AA) (**Figure 1A**), and containing an N-terminal signal peptide for 293/357 (80%) of them, as predicted by SignalP5.0 (**Supplementary Table 4**). Overall, 212 (∼60%) did not contain the known specific effector features predicted by EFFECTORP-fungi 3.0 (**Figure 1B, Supplementary Table 5**), suggesting that they may represent novel structural features not yet implemented in the EFFECTORP-fungi 3.0 model. At the primary sequence level, 187 were unique, while 170 shared similarities with another member of the 357 core secretory proteins (**Supplementary Table 6)**. We also observed that while 113 did not yield any functional annotation, 244 (68%) harboured conserved domains referenced in the pfam database (**Figure 1C**, **Supplementary Table 7**).

**Figure 1.**
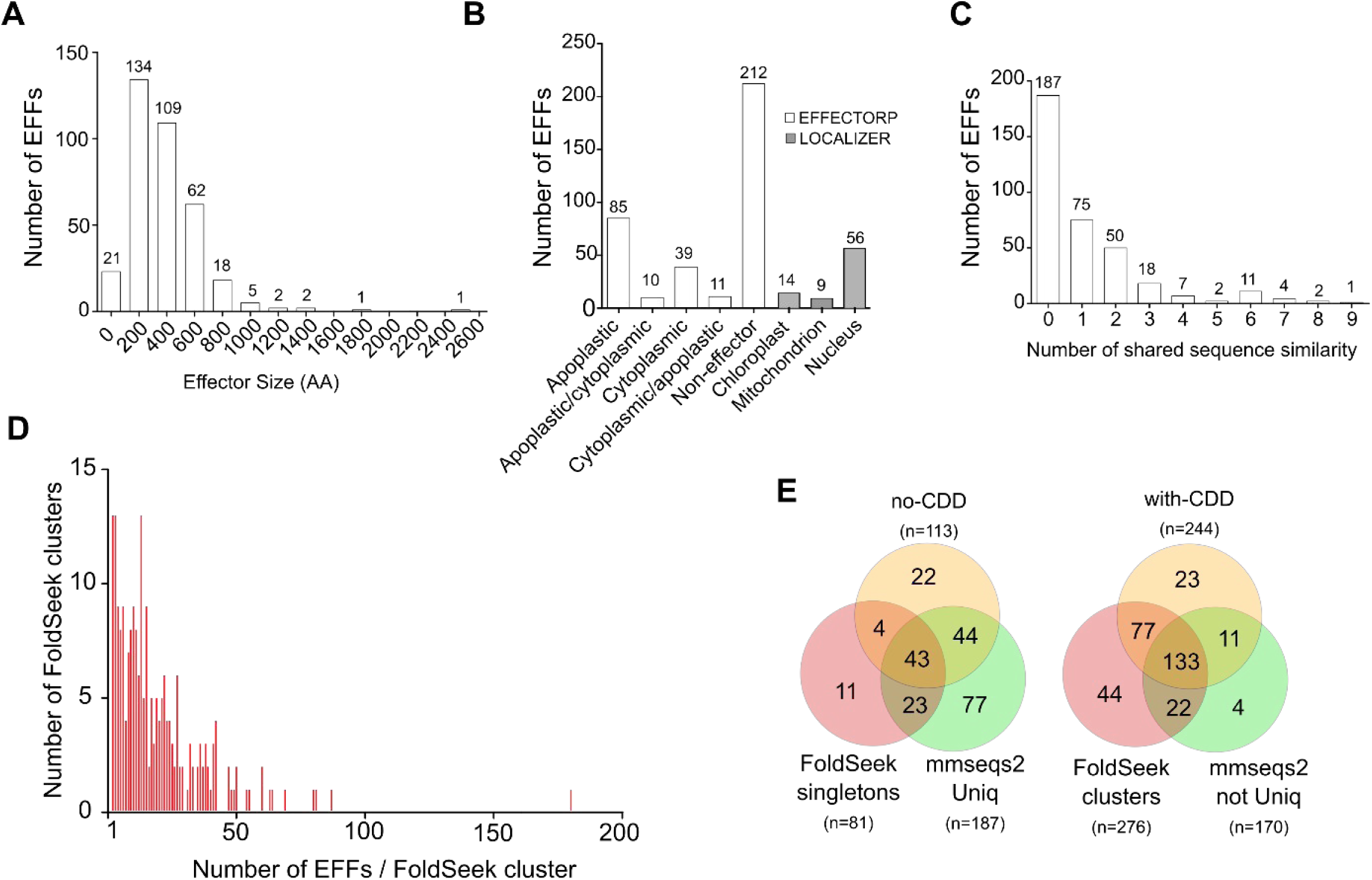
Characteristics of the 357 core secretory proteins. Classification of the 357 core secretory proteins according to A) their size in amino acids (AA), B) EFFECTORP-fungi 3.0 and LOCALIZER1.0.4 predictions and C) protein sequence similarities shared within the 357 secretory proteins (EFFs). D) FoldSeek clustering based on structural similarities between the 357 EFFs and putative effectors described in three recent publications (Rocafort et al., 2022; Derbyshire and Raffaele, 2023 and Seong and Krasileva, 2023). Identified clusters are represented according to their sizes ranging from 2 to 180 (the 81 singletons are not shown). E) Venn diagrams representing the unique (left) and common (right) protein features cross-referencing (i) shared sequence similarities identified by MMSEQS2, (ii) shared conserved domains referenced in pfam predicted by a Conserved Domain Database (CDD) analysis and (iii) shared structural similarities defined by FoldSeek clustering.

We then used FoldSeek (van Kempen *et al*., 2023) to cluster the 357 proteins according their protein structure similarities. First, predicted structures were successfully collected for 351/357 and looking at their pLDDT (Predicted Local Distance Difference Test) used as a metric to assess the reliability of predictions (**Supplementary Table 8)**, 92/351 have pLDDT below 0.7 and 26/351 below 0.5. While low values may impair the clustering efficiency, 47 clusters and 245 singletons (∼70%) were detected, depicting substantial heterogeneity in their structural folds. Among evidenced identified clusters, 39, 5, 2 and 1 gathered 2, 3, 4 or 5 secretory proteins, respectively (**Supplementary Table 8)**. In a second strategy, clustering was performed between the 351 core secretory proteins supplemented with 3,991 known secreted proteins identified in three recent studies from different pathogens (Rocafort *et al*., 2022; Derbyshire and Raffaele, 2023; Seong and Krasileva 2023). This analysis narrowed down to 81 singletons (22%), while the remaining proteins were grouped into clusters of varying sizes, including a large cluster of 180 members and 52 others belonging to clusters containing 2 to 87 members (**Figure 1D**, **Supplementary Table 9**). Comparing the different approaches of domain analysis, similarity searches between the protein sequences of the 357 proteins and their clustering with other published fungi secretomes, 43 proteins (12% of our 351 core secretome) remained unannotated, highlighting the distance still to be covered to fully characterize the *F. graminearum* secretome, while 113 (32%) contained protein domains shared between different members of the secretome and were included in FoldSeek clusters already described (**Figure 1E**, **Supplementary Table 8** and **9**). These preliminary analyses suggest that the 357 core secretory proteins represent an original source of unexplored and uncharacterized effectors with pivotal role during the early stages of the FHB disease.

### Identification of secretory proteins targeting the plant nucleus

As already suggested by Rocher *et al*. (2022), we hypothesized that a subset of the secretory proteins could be imported into the plant nucleus. The predicted localization of the 357 secretory proteins using the current version of the LOCALIZER tool identified 56 secretory protein sequences putatively able to enter the plant nucleus (**Figure 1B** and **Supplementary Table 10)**. Among these, 14 core secretory proteins were further short listed based both on their consistent expression changes during infection in different strains and on their expression profiles that matched with the effector delivery waves hypothesis (Fabre *et al*., 2019; Rocher *et al*., 2022) (**Table 1)**. More specifically, this subset included six and eight secretory proteins belonging to the early and late delivery wave, respectively, with fold changes ranging from 0.48 to 4.55 Log_2_ normalized counts between 48 and 72 hours post inoculation (hpi) (Rocher *et al*., 2022).

**Table 1.** Selected set of 14 secretory proteins predicted by LOCALIZER1.0.4 to be targeted to the plant nucleus. Molecular features of 14 selected proteins expected to be targeted to the nucleus by LOCALIZER1.04 and characterized using EFFECTORP3.0, SIGNALP5.0, their expression profiles and Log2 fold change (FC) during wheat spike infection from Rocher et al 2022.

| Accession ID | Code | SIZE (AA) | Effector Prediction (EFFECTORP) | Signal Peptide Prediction (SIGNALP) | Nuclear Localization Prediction (LOCALIZER) | Expression pattern during FHB | Log(FC) between 48hpi and 72hpi | Fast and slow evolving genome fraction |
| --- | --- | --- | --- | --- | --- | --- | --- | --- |
| FGRAMPH1_01T01843 | EFF01 | 383 | Non-effector | yes | Nucleus | Late | 0.54 | Fast |
| FGRAMPH1_01T05171 | EFF02 | 617 | Non-effector | yes | Nucleus | Late | 1.17 | Fast |
| FGRAMPH1_01T11497 | EFF03 | 249 | Cytoplasmic effector | yes | Nucleus | Early | 4.55 | Fast |
| FGRAMPH1_01T11559 | EFF04 | 247 | Cytoplasmic effector | yes | Nucleus | Late | 0.49 | Fast |
| FGRAMPH1_01T11579 | EFF06 | 856 | Non-effector | yes | Nucleus | Late | 2.25 | Fast |
| FGRAMPH1_01T12575 | EFF10 | 1009 | Non-effector | yes | Nucleus | Early | 0.54 | Fast |
| FGRAMPH1_01T13187 | EFF11 | 528 | Non-effector | yes | Nucleus | Early | 0.82 | Fast |
| FGRAMPH1_01T13647 | EFF13 | 663 | Non-effector | yes | Nucleus, Apoplast | Early | 1.26 | Fast |
| FGRAMPH1_01T13999 | EFF17 | 1350 | Non-effector | yes | Nucleus, chloroplast | Early | 0.76 | Fast |
| FGRAMPH1_01T15319 | EFF18 | 439 | Non-effector | yes | Nucleus | Late | 1.25 | Fast |
| FGRAMPH1_01T18783 | EFF20 | 569 | Non-effector | yes | Nucleus | Early | 2.42 | Fast |
| FGRAMPH1_01T21315 | EFF22 | 441 | Cytoplasmic effector | yes | Nucleus | Late | 0.86 | Slow |
| FGRAMPH1_01T21741 | EFF24 | 876 | Non-effector | yes | Nucleus | Late | 0.50 | Slow |
| FGRAMPH1_01T25857 | EFF29 | 505 | Non-effector | yes | Nucleus, Apoplast | Late | 3.94 | Fast |

To confirm the nuclear localization of the 14 selected secretory proteins *in planta*, GFP-fused constructs without signal peptide were designed for further Agrobacterium-mediated transient expression in *N. benthamiana* leaf cells using an NLS-mRFP marker as a control. Confocal microscopy observations from at least two independent experiments revealed that nine out of 14 proteins effectively accumulated in the nucleus (**Figure 2A**). The remaining five proteins, which either showed weak expression or did not localize to the nucleus, such as EFF02 and EFF24 (**Figure 2A**) served as valuable negative controls for this experiment, demonstrating the specificity of our nuclear localization assay. Among the nine nuclear localized proteins, distinct sub-nuclear localizations were observed. For instance, EFF18-GFP fusion protein showed a strong nucleoplasm accumulation and was excluded from the nucleolus while EFF01-GFP strongly accumulated into the nucleolus. Although the observations were reproducible, some secretory protein-GFP fusion proteins, such as EFF20-GFP, EFF22-GFP and EFF29-GFP, consistently exhibited weak fluorescence signals or display distinct nuclear localization as EFF03-GFP being either nucleolar or nucleoplasmic (**Figure 2A**).

**Figure 2.**
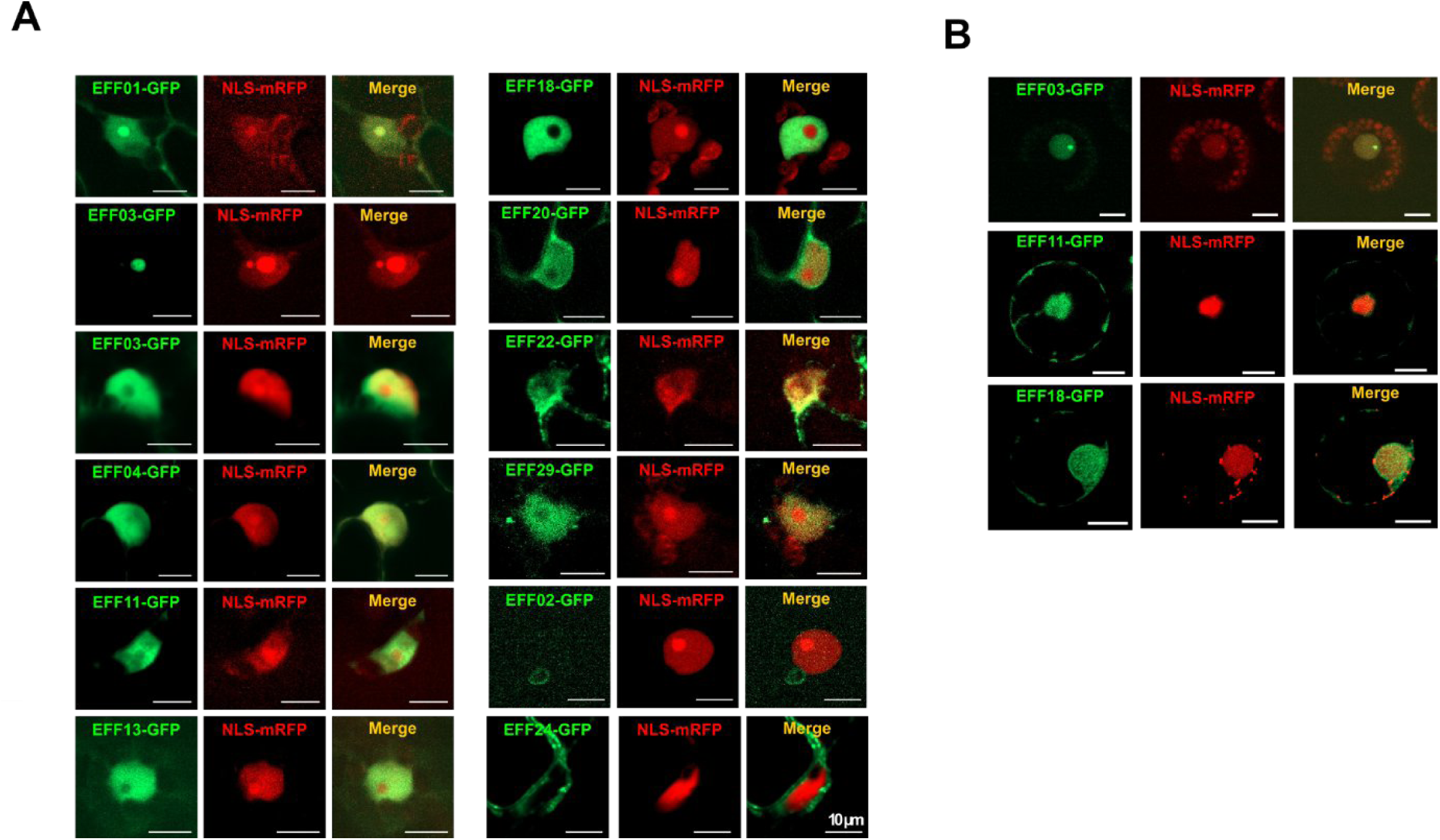
Characterization of 14 secretory proteins by transient expression in *Nicotiana benthamiana* and *Triticum aestivum*. **A-B)** Localization of co-transfected EFF-GFP and NLS-mRFP in transiently expressed *N. benthamiana* (**A**) or in wheat protoplasts (**B**) recorded three days after the inoculation respectively by spinning disk microscopy under 40X objective and confocal microscopy under a 63X oil objective. EFF02 and EFF24 that are not addressed to the nucleus are used as negative controls.

To further demonstrate that secretory proteins effectively target the nucleus of the host cells, transient expression in wheat protoplasts was repeated for a subset of three proteins that showed strong nuclear accumulation in *N. benthamiana*, but with distinct subnuclear localization (EFF03-GFP, EFF11-GFP and EFF18-GFP). All three secretory proteins-GFP fusion proved to also accumulate in wheat nuclei. Similar sub-nuclear localization was observed for EFF11-GFP fusion proteins in *N. benthamiana* and wheat, while the fluorescence signal of EFF03-GFP was mainly nucleolar in wheat and the localization of the EFF18-GFP fusion protein was heterogeneous and appeared in small domains throughout the wheat nucleus as well as potentially within the nuclear membrane, as observed in the co-localization experiment with NLS-mRFP (**Figure 2B**).

### Molecular characterization of the nuclear-targeted secretory proteins

To get further insights into the nine selected secretory proteins, which localize to the plant nucleus, we then explored their molecular structure using a set of *in silico* tools. First, conserved domains were defined using the Conserved Domain Database (CDD) (Lu *et al*., 2020). Despite the small size of the core secretory proteins, most of them, except EFF03, contained one or more protein domains referenced in the pfam database, with EFF18 and EFF22 sharing a common pfam domain (peptidase_M14; pfam00246) (**Figure 3A**). Many of these domains were annotated as peptidases and hydrolases, suggesting a function in protein degradation and catabolism, while others have possible functions in the redox metabolism **(Supplementary Table 6)**. Amino acid identity of the nine nuclear-targeted proteins were searched among a set of 22 fungal species whose genomes are fully sequenced, including several *Fusarium* species up to *Ustilago maydis*, a more distant species. Seven proteins were found in 21 to 22 fungal species and the mean percentage of identity of the nine proteins was of 73.9±23.9% (mean without *Fgr*). This suggest that the nine core secretory proteins are well-conserved within the Ascomycota division when compared to the well conserved RPA protein (82.5±18.5) or a tyrosine kinase protein (52.8±29.0) selected for its high accumulation of non-synonymous mutations (**Figure 3B** and **Supplementary Table 11**). Interestingly, while EFF18 and EFF22 shared a common pfam domain, they do not belong to the same FoldSeek cluster (**Supplementary table 7** and **8**), indicating that secondary and tertiary structural predictions provide complementary information. This prompted us to study the tertiary structures of the nine secretory proteins using ColabFold (**Figure 3C**). Most proteins displayed a very compact globular structure made of helixes and beta sheets, while EFF03 and EFF04 are not well predicted with a high Predicted Template Modeling (pTM) average value of 0.32 and 0.61 respectively. Indeed, low percentages of disorder (absence of protein structure) were predicted in the nine proteins (mean: 16±11%, **Supplementary Table 12**) using DEPICTOR2 (Basu *et al*., 2023). Again, among the nine core secretory proteins, EFF03 and EFF04 are exceptions with 47% and 29% of disorder, respectively (**Figure 3D**) which corroborate their low pTM values. Finally, despite their high level of identity across a large panel of fungi species, eight out of nine proteins belonged to the fast-evolving fraction of the *Fusarium* genome (**Figure 3E**). Taken together, the nine secretory proteins form highly compact tertiary structures and as core secretory proteins, they were conserved among a large panel of fungal species.

**Figure 3.**
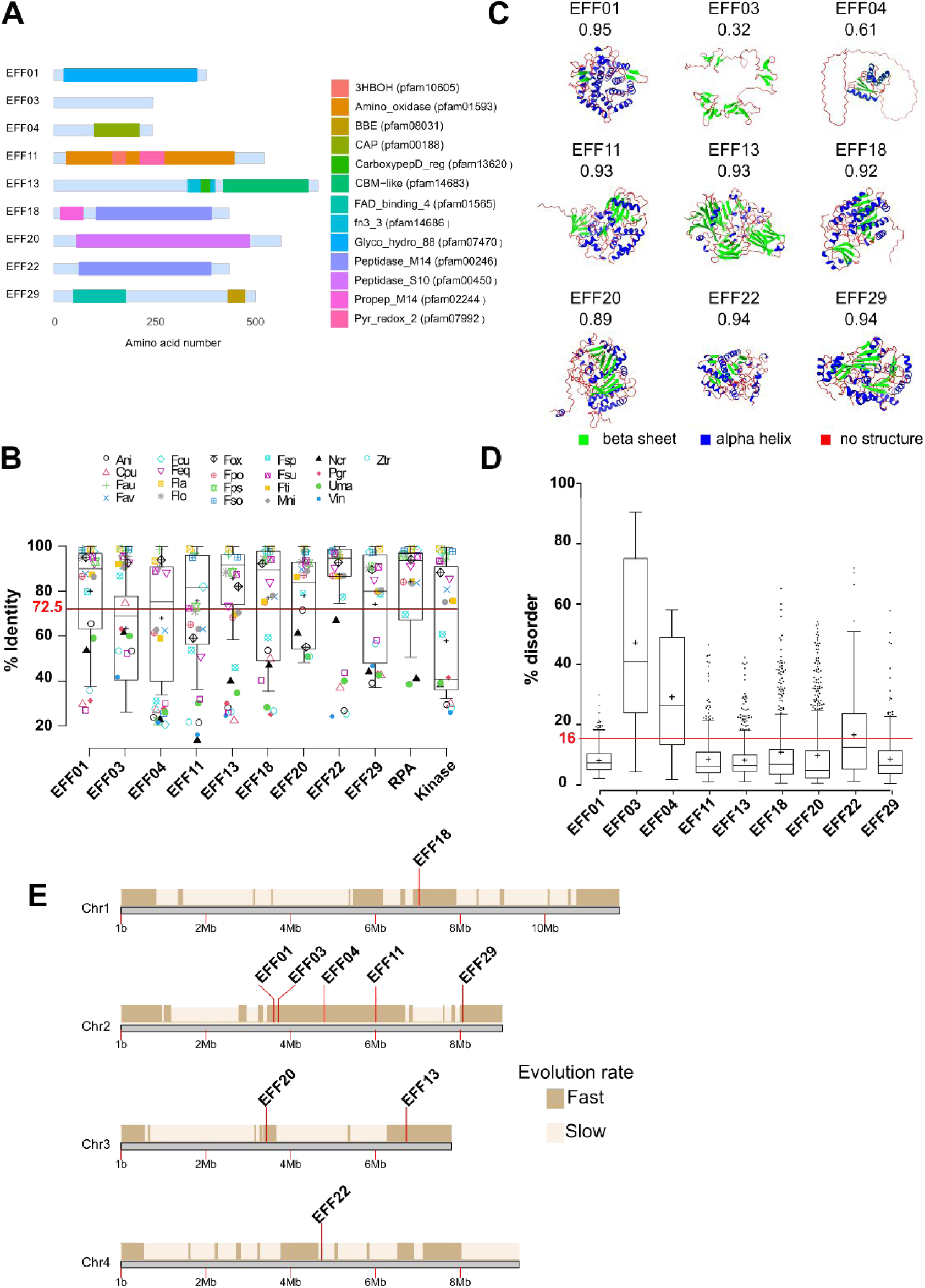
Characteristics of 9 secreted proteins from *F. graminearum* targeted to the plant nucleus. A) Conserved secondary structures of the 9 secreted proteins as predicted by the Conserved Domain Database (CDD) analysis. B) Box plots with Tukey whiskers representing the percentage of the 9 secretory protein identity (% identity) in 22 species of fungi as well as for RPA and a tyrosine kinase protein (kinase) chosen for their low and high conservation, respectively. Mean value (72.5%) of percentage of identity of the 9 secreted proteins is indicated as a red line. C) 3D structures predicted by ColabFold and refined by ChimeraX to highlight beta-sheets (green) and alpha helixes (blue) secondary structures. pTM values ranging from 0 to 1, with higher scores indicating better structural predictions are indicated below each structure. D) Box plots with Tukey whiskers representing the percentage of intrinsically disordered regions (% disorder) of the 9 selected proteins. Mean value (16%) of disorder of the 9 secretory proteins is indicated as a red line. E) Positions of the nine genes encoding the secretory proteins according to the fast and slow evolving regions along the 4 chromosomes of *F. graminearum* genome.

### Identification of wheat proteins interacting with EFF18

To identify potential interactors of the secretory proteins that may act as susceptibility genes, we employed the yeast two-hybrid (Y2H) assay, a classical method for detecting direct protein-protein interactions. We selected EFF18 as a candidate to perform a Y2H screen, as EFF18 was consistently expressed in host plant with efficient transient expression assay that undoubtedly confirmed its nuclear localization in wheat protoplasts. EFF18 is well conserved among 22 pathogen species, contains low disorder, high pTM (**Figure 2C-D**) and its expression varies significantly (**Table 1**) between 48 and 72 hpi. To optimize our chance to identify relevant protein interactors, a specific cDNA library was generated using the wheat Recital cultivar infected with the MDC_Fg1 strain, as in Rocher *et al*. (2022).

A first Y2H screen was carried out using the stringent LexA system yielding 20 positive clones, among which only one cDNA sequence (LOC123138250), annotated as a 5-oxoprolinase-like, was identified as multiple clones with a high confidence B score. A second screen, carried out with the more sensitive Gal4 system, identified 100 positive clones, four of them having high confidence scores (A or B). As in the LexA screen, LOC123138250 was identified, as well as three more cDNAs: LOC123105923, encoding a beta-amylase 2 chloroplastic-like protein involved in polysaccharide catabolic processes, LOC543119 annotated as thylakoid formation 1 involved in the photosystem II assembly in chloroplasts and LOC123116977, an ER membrane protein complex subunit 1-like acting in the glutathione metabolic process (**Supplementary Table 13**). The four proteins included conserved domains referenced in pfam (**Figure 4A**). Consistent with EFF18’s classification as a member of the core secretome targeting conserved function in the host, we found that all four targets were conserved across a large panel of plant species representing the green lineage, exhibiting an average sequence identity of 67±10.8% (**Supplementary Table 14**). This value positioned the four targets as well-conserved proteins when compared to the MUTS HOMOLOG 1 (MSH1), a miss-match repair protein involved in genome stability which is well-conserved in the green lineage (Bai and Guo, 2023) and to the AGAMOUS-LIKE 62 (AGL62) protein, a member of type I MADS-Box transcription factors described to have a high rate of gene duplication and reduced purifying selection (Nam *et al*., 2004) with a percentage of identity of 61.1 ±10.8% and 36.9±5% respectively (**Supplementary Table 14**). The phylogenetic trees illustrate this conservation with evolution rates expressed as the sum of the branch lengths for monocot species, similar to MSH1 but lower than AGL62 (**Figure 4B**). In good agreement with the fact that the Y2H library was constructed from spikes tissues, analysis of previous transcriptomic data gained in wheat spikes between 48 and 96 hpi (Rocher et al., 2024) indicated that the four proteins are expressed in spikes with a dynamic pattern of expression during infection. LOC543119 and LOC123105923 are induced in response to Fusarium infection, with mRNA levels increasing as infection progresses. Instead, LOC123116977 transcript levels decrease in response to infection, while expression of the LOC123138250 is not affected by *F. graminearum* infection (**Figure 4C** and **supplementary Table 15**). Interactions between EFF18 and its four putative interactors were then confirmed by performing an Y2H interaction assay carried out by reversing the bait and the prey (**Figure 4D)**. Finally, to determine whether the four targets were addressed to the nucleus as for EFF18, a transient expression assay was performed revealing that all four proteins were effectively addressed to and accumulated in the nucleus when co-expressed with EFF18 in *N. benthamiana* (**Figure 4E**), suggesting that they could interact physically within the nucleus. In this assay, we also observed that EFF18 formed aggregates as observed in the transient expression in wheat protoplasts (**Figure 2B**) when co-transfected with LOC123116977 and LOC123105923. Taking together, these results suggest that EFF18 and its four interacting targets are conserved across a large panel of species, are expressed in spikes after anthesis during pivotal stages of the FHB progress and that both EFF18 and its targets can be concurrently addressed to the plant nucleus.

**Figure 4.**
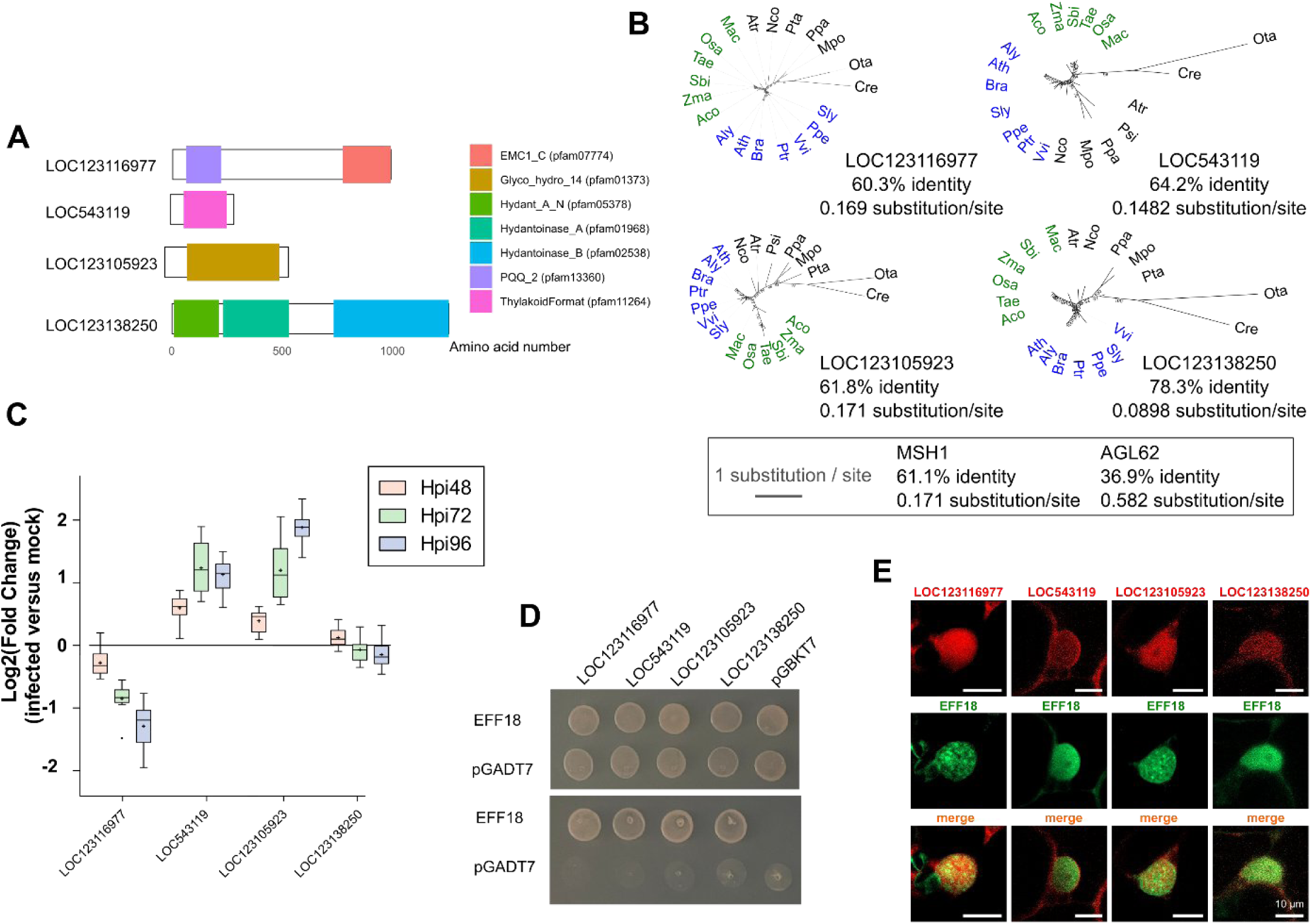
Characteristics of EFF18 target genes identified by Y2H screen. **A)** Protein domains identified in the 4 target proteins as predicted by the Conserved Domain Database (CDD) analysis. **B**) Unrooted phylogenetic trees where monocotyledons (green) and dicotyledons (blue) are indicated. Branch lengths were standardized for all trees and expressed as substitution/site. Percentage of identity (%identity) and evolution rates computed as the sum of the branch lengths for the monocotyledons species are given below each tree. Values for AGL62 and MSH1 served as indicators for high and low evolution rates. **C)** Expression profiles of the 4 target proteins in the ‘Recital’ wheat cultivar. Data are expressed as Log2(FC) between infected (values obtained from plants facing 3 different *F. graminearum* strains in 3 replicates each) and mock (3 replicates) conditions at 48, 72 and 96 hours after infection (Hpi). **D)** Y2H interaction assay between EFF18 (prey) and its four interacting proteins (baits). Y2H assay was performed using permissive (YNB/-Leu/-Trp, top panel) or test (YNB/-Leu/-Trp/-Ade/-His; bottom panel) media. Empty vector pGADT7 and pGBKT7 were used respectively as prey and bait negative controls. **E)** Co-localization assay of the four target proteins expressed as mRFP fusions and EFF18-GFP in transient expression assay in *N. benthamiana* after three days observed using a Zeiss Cell Observer Spinning Disk microscope under 40X objective. Scale bar: 10µm.

### Characterization of the interactions between EFF18 and the beta-amylase target

ColabFold_Multimer tool was used to predict the 3D structures of the complexes between EFF18 and its four interacting targets. The Predicted Aligned Error (PAE) score that defines the confidence of the residue positions within the predicted structures showed that intra- and intermolecular interactions were predicted (**Figure 5A**). However, for all four predicted complexes, evaluation of the quality of interactions between residues by the interface predicted Template Modeling (ipTM) score, gave low scores, below 0.7, that is commonly used as a threshold to predict significant protein-protein interactions. This is likely explained by the low predicted Template Modeling (pTM) score ranging from 0.59 to 0.74 (**Figure 5A**). For this reason and to avoid considering the protein complex as a whole, local interactions were predicted by using Alphafold-Multimer Local Interaction Score (AFM-LIS) (Kim *et al*., 2024). AFM-LIS predicted the interaction between EFF18 and LOC123105923 (beta-amylase 2), but failed to detect any other interaction (**Table 2**). The interacting residues at a distance of less than 5 Ångströms from each other were then predicted with ChimeraX (**Figure 5B, 5C** and **Supplementary Table 16)**. This analysis identifies one main region in EFF18 between position 304 to 350AA (45 AA), including a large alpha helix (313-330 AA), and four smaller interacting regions in the LOC123105923 between 226 to 422AA (196AA) (**Figure 5D**). This prediction was used to design deletion derivatives of EFF18, in which 45 AA or its C-terminal region was removed (**Figure 5E**). Using these new constructs in an Y2H interacting assay, we demonstrated that deleting the 45AA region was sufficient to impair the interaction between EFF18 and the LOC123105923 (**Figure 5F**), without affecting the predicted structure of EFF18 (**Supplementary Figure 1**). To sum-up, the ColabFold_Multimer tool predicted complexes with low ipTM scores (<0.7) indicating weak protein-protein interactions. Prompting the use of AFM-LIS to confirm interaction with LOC123105923, further validated the interaction by identifying key interacting residues and regions using ChimeraX and functional assay in Y2H.

**Figure 5.**
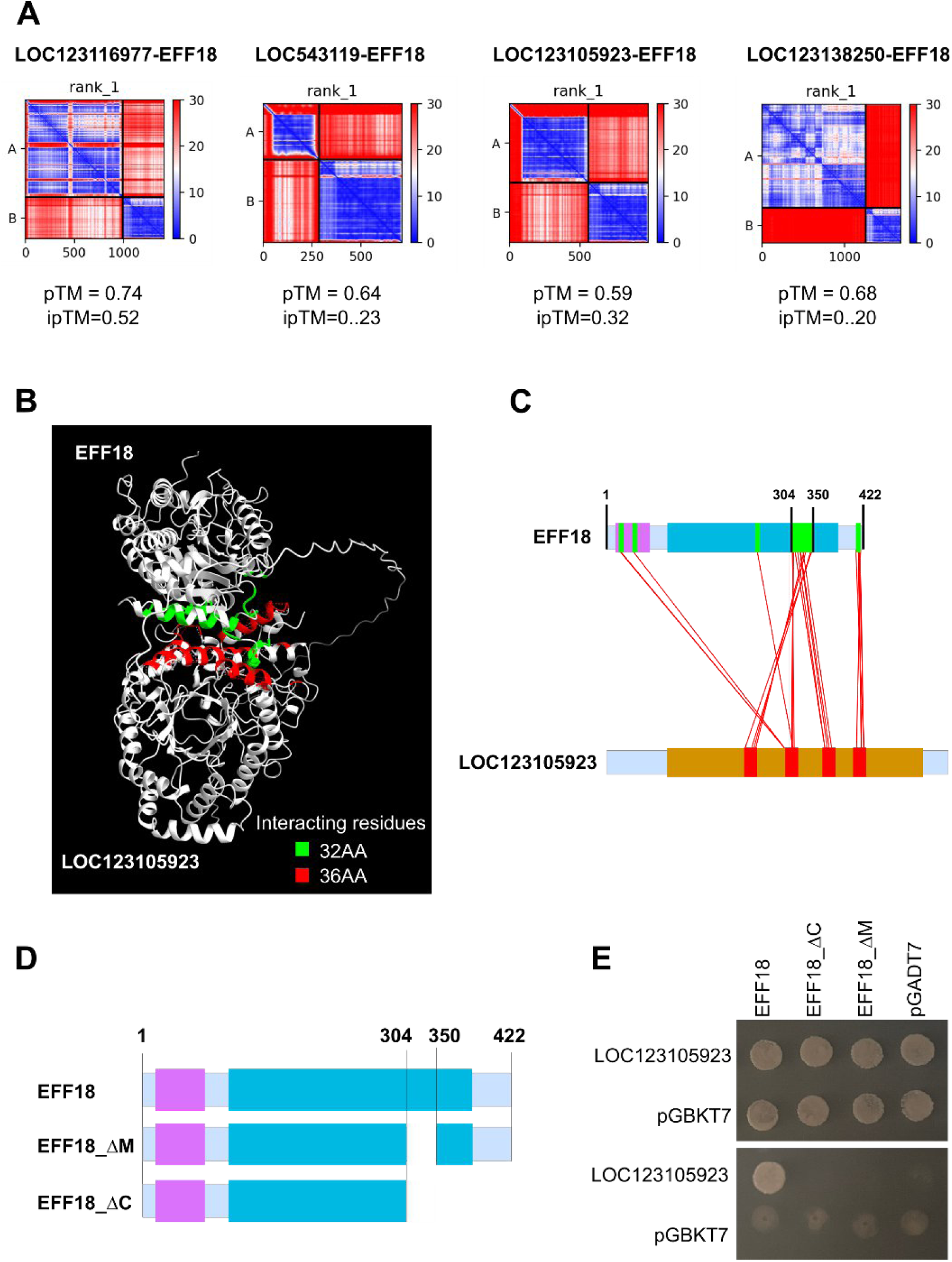
Interactions between EFF18 and its four targets predicted by ColabFold_multimer. **A)** Predicted Aligned Error (PAE) score obtained from a ColabFold_multimer prediction and representing the probability of amino acids interacting with each other. PAE plots are represented as 2D matrixes with protein A as the targets and B as EFF18. PAE values range from 0 to 30 Ångströms (y-axis), where 0 indicates high (blue) and 30 low (red) confidence in interactions, along the protein (x-axis in amino acids). Intra and intermolecular interactions can be observed for A top left, B bottom right and A-B top right, B-A bottom respectively. pTM and ipTM of the complexes are indicated below the corresponding PAE plots. **B)** Protein complex with interacting residues highlighted in green (EFF18, 32AA, top) and red (LOC123105923, 36AA, bottom). **C)** Definition of the interacting residues between EFF18 and LOC123105923 using ChimeraX and redrawn at scale highlighting EFF18 (green) and beta amylase (red) residues in interaction. Pfam domains identified by the Conserved Domain Database (CDD) are indicated as in Figure 4. **D)** EFF18 deletion derivatives used in Y2H assay. **E)** Y2H assay between EFF18, EFF18 deletion derivatives and LOC123105923. The prey (in pGADT7 vector) together with each of the bait plasmids (in pGBKT7 vector) were transformed into the yeast strains Y187 and AH109 respectively. Y2H assay was performed using permissive (YNB/-Leu/-Trp, top panel) or test (YNB/-Leu/-Trp/-Ade/-His, bottom panel) media. Empty vector pGADT7 and pGBKT7 were used respectively as pray and bait negative controls.

**Table 2.**
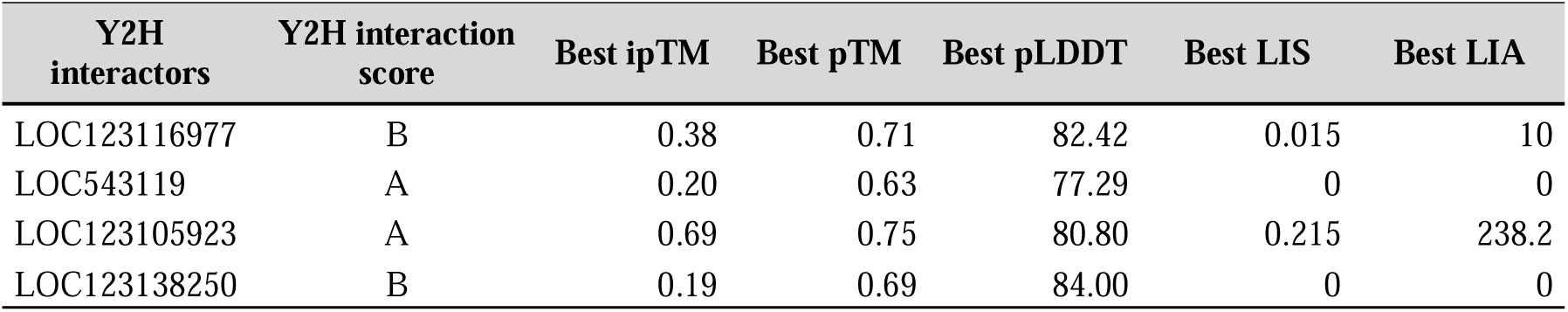
Interaction scores for 4 complexes between targets and EFF18 in Y2H and with AFM-LIS. From the PAE analysis (probability of interaction between amino acids based on the distance in Ångströms between 2 amino acids), AFM-LIS allows predicting PPIs at the scale of each amino acid by jointly using a Local Interaction Area (LIA) and a Local Interaction Score (LIS). AFM-LIS with best Local Interaction Area (LIA) and best Local Interaction Score (LIS) above the minimal values (respectively LIA ≥ 3432 and LIS ≥ 0.203.

## Discussion

In this study, we examined putative core secretory proteins expressed by several strains of *F. graminearum* of varying aggressiveness in different wheat cultivars during FHB infection and predicted to be targeted to the plant nucleus. These effector-like proteins were found to be significantly regulated during the FHB infection dynamics, suggesting a pivotal role in the infection success. Among the 357 putative secretory proteins described in our previous study (Rocher *et al*., 2022), 56 were predicted to be localized in the nucleus. In this study, we shortlisted 14 nuclear secretory proteins, all of which harbored a predicted nuclear localization signal (NLS). Transient expression of these proteins in *N. benthamiana* demonstrated the effective nuclear localization for more than two third with specific subnuclear accumulations, and three were further successfully validated in wheat cell nuclei. By targeting a specific subnuclear structure, these effector-like proteins may interfere with various critical host processes, exerting broad-spectrum effects on the primary functions of the plant cells, thereby helping the pathogen in establishing, proliferating and spreading in infected host tissues. Several nuclear-targeting effectors have already been described in plant pathogens, such as Mlp124478 from *Melampsora larici-populina* known to disrupt host gene expression by directly interacting with plant transcription factors (Petre *et al*., 2016) and VdSCP7 from *Verticilium dahliae* that targets the plant nucleus and alters plant immunity (Zhang *et al*., 2017). In *F. graminearum*, very few nuclear-targeting effectors are known. FgNLS1 has been recently described for its ability to reach the cell nuclei and to suppress plant immunity at early stages of the infection (Hao *et al*., 2023). Here, this study suggests that *F. graminearum* likely deploys a complex arsenal of effector-like molecules into wheat cell nuclei, enabling multiple nuclear targets to manipulate complementary host functions effectively. Interestingly, in addition to the strong sequence conservation and expression observed among *F. graminearum* strains with contrasting levels of aggressiveness, 7 out of the 9 nuclear-targeting secretory proteins were also well conserved across 22 fungal species, as exemplified by the high conservation of EFF18 in our phylogenetic analysis. This suggests that these secretory proteins may serve as generic components of infection mechanisms shared by multiple pathogens across diverse host plants. Such conserved drivers of pathogenicity among various infectious pathogens supports the idea that common functions, likely essential for the infectious process, are maintained across species to facilitate infection in diverse host plants. Examples of conserved nuclear effectors playing a significant role in the infection process by manipulating primary host defense responses have already been reported. For example, XoPD, a type III secretion effector from *Xanthomonas euvesicatoria*, plays an important role in altering host gene expression, a common strategy among different pathogens (Kim *et al*., 2013). By targeting host transcription factors such as SIERF4, this effector is able to suppress ethylene production, thereby further reducing defense responses. Similarly, the Transcription activator-like effectors (TALEs) are important effectors found in several pathogenic bacteria, such as *Ralstonia solanacearum* and *Xanthomonas* spp, able to act as specific plant transcription factors via a programmable DNA-binding domain (Hutin *et al*., 2015). TALEs from different bacterial strains of the same pathogen have been showed to target the same susceptibility genes and thus have conserved interactors (Boch *et al*., 2014). Taken together, the nuclear localization and broad conservation of these proteins across diverse pathogens underscore their potential role as nuclear core effector proteins able to promote generic determinants of pathogenicity across a wide range of crops. Because nuclear effectors are supposed to interact directly or indirectly with host susceptibility factors by targeting specific players of the host cellular processes, we chose EFF18 as a case study to identify new drivers of FHB susceptibility in wheat. EFF18 was chosen because of its large accumulation in the nucleus and of its increasing expression patterns at the early stages of the FHB infection (Rocher *et al*., 2022). We then screened for novel interacting partners using a Y2H assay performed on an original and relevant cDNA library obtained from wheat spikes infected with *F. graminearum*. Using this strategy, we identified several plant proteins, which may represent new susceptibility genes able to control FHB development in wheat spikelets. LOC123105923 is a predicted beta-amylase protein involved in the hydrolysis of starch to maltose. In Sweet potato tubers, it plays an important role in carbohydrate metabolism by converting stored starch into sugar, which provides energy for many metabolic processes, including growth and development (Vajravijayan *et al*., 2018). LOC543119 is involved in the formation of thylakoids in chloroplasts and is essential for photosynthesis. Its role in photosynthesis and further in plant health and defense has already been reported (Monroe and Storm, 2018). LOC123116977 is an endoplasmic reticulum (ER) membrane protein and is important for various cellular processes including stress response including defense response (Yamada *et al*., 2011) and, LOC123138250 is a 5-oxoprolinase that has also been observed to be involved in stress response (Lei *et al*., 2024). While initially not predicted in the nucleus, we find them to localize to the nucleus in presence of the EFF18 in transient expression assays, suggesting that they may have additional nuclear functions than the ones predicted or demonstrated so far. Only the EFF18/LOC123105923 interaction was successfully confirmed by ColabFold predictions motivating further attention to these interacting candidates. Interestingly, a plant beta-amylase proved to accumulate in a susceptible triticale line after infection but not in resistant ones, suggesting that inhibition of alpha- and beta-amylase activities from pathogen may be associated with Type IV resistance to FHB (Perlikowski *et al*., 2016). Diverting sugar metabolism is a usual effector target for plant pathogens to create favorable conditions for their growth and spreading (Vajravijayan *et al*., 2018). This beta-amylase was highly conserved across the green lineage, echoing with the high sequence conservation of its bait, EFF18, and emphasizing that relative pathogens could target conserved processes across crops. Although beta-amylases are not predicted to accumulate in the nucleus, the effective nuclear colocalisation of LOC123105923 with EFF18 was clearly proved in this study, strengthening their value for further analyses to understand the beta-amylase role in plant susceptibility to pathogens.

To conclude, parsing multiple candidate *F. graminearum* genes supposed to control FHB in wheat, this study provides new insights about the sophisticated strategy used by this pathogenic fungus, emphasizing its ability to deploy a complex arsenal of effector-like proteins towards multiple sub-nuclear wheat targets at early stages of infection to manipulate complementary host functions. Being part of evolutionarily conserved elements across diverse pathogens, these fungal proteins could be at the crossroad of shared mechanisms of pathogenicity that could facilitate infection in a broad range of host plants through the control of generic susceptibility pathways. While further investigations are required, a more in-depth characterization of such fungal core secretory proteins and their plant targets could unlock new opportunities for identifying novel resistance sources, enabling the development of robust, broad-spectrum strategies to control multiple pathogens across various crops.

## Supplementary data

**Supplemental Table 1** provides the list of plant species used for similarity search using protein sequences and phylogenetic analyses.

**Supplemental Table 2** provides the list of fungi species used for similarity search using protein sequences and phylogenetic analyses.

**Supplementary Table 3** provides the list of 357 core secretory proteins defined by Rocher et al 2022 and their codification as “EFF” used in this article.

**Supplementary Table 4** provides SignalP5.0 prediction of the occurrence of Signal peptide for of the 357 core secretory proteins defined by Rocher *et al*. 2022.

**Supplementary Table 5** provides EFFECTORP3.0 prediction using protein features usually found in know effectors for the 357 core secretory proteins defined by Rocher *et al*. 2022.

**Supplementary Table 6** provides unique or shared features of the 357 core secretory proteins defined by Rocher *et al*. 2022 and used to construct Venn diagrams.

**Supplemental Table 7** provides Conserved Domain Database (CDD) analysis to identify pfam protein domains in the 357 core secretory proteins defined by Rocher *et al*. 2022.

**Supplemental Table 8** provides a first FoldSeek clustering results of 351 core secretory proteins defined by Rocher *et al*. 2022.

**Supplementary Table 9** provides a second FoldSeek clustering results of the 351 core secretory proteins defined by Rocher *et al*. 2022 with putative effectors from the literature available in an in-house database.

**Supplementary Table 10** provides a LOCALIZER1.0.4 prediction of the 357 core secretory proteins defined by Rocher *et al*. 2022.

**Supplementary Table 11** provides percentage of protein identity for the 9 proteins localized into the nucleus in transient expression.

**Supplementary Table 12** provides a prediction of protein disorder for the 9 proteins localized into the nucleus in transient expression.

**Supplementary Table 13** provides the raw data provide by the Hybrigenics company for the identification of interacting partners for EFF18 by yeast-2-hybrid screens.

**Supplementary Table 14** provides percentage of protein identity in the plant lineage of the 4 interacting proteins with EFF18 selected from Y2H screens.

**Supplementary Table 15** provides expression profiles of the 4 interacting targets during wheat spike infection.

**Supplementary Table16** provides prediction of interacting residues <5 Angstroms by ChimeraX.

**Supplementary Figure 1** shows the structural conservation between EFF18 and EFF18_ΔM

## Acknowledgments

C.T. and L.B. would like to thank Dr Freddy Boutrot and Dr Aline Probst for critical review of the manuscript. The authors thank Richard Blanc for its valuable help during plant experiments and *Fusarium graminearum* inoculations. All microscopic images were acquired on the CLIC microscopy facility (CLermont Imagerie Confocale). All images were stored in an OMERO server hosted at the Mesocentre Clermont Auvergne. The authors thank Richard Blanc for its valuable help during plant experiments and *Fusarium graminearum* inoculations.

## Author contributions

L.B. and C.T. supervised the study. S.A., E.V., L.L., F.R., and M.B. performed experiments. S.A., C.T. and M.B. performed the bio-informatics work. S.A., L.B., and C.T. wrote the manuscript.

## Data availability

All data are provided in Supplementary files.

## Abbreviations

AFM-LIS: Alphafold-Multimer Local Interaction Score
AGL62: AGAMOUS-LIKE 62
ADP-glucose pyrophosphorylase: AGPase
CDD: Conserved Domain Database
*F. graminearum*: *Fusarium graminearum*
FHB: Fusarium head blight
GFP: Green Fluorecent Protein
hpi: hours post inoculation
ipTM: Interface Predicted Template Modeling
LIA: Local Interaction Area
LIS: Local Interaction Score
mRFP: Monomeric red fluorescent protein
MSH1: MUTS HOMOLOG 1
*N. benthamiana*: *Nicotiana benthamiana*
NLS: Nuclear Localization Signals
PBS: Possible Biological Score
PAE: Predicted Aligned Error
pTM: Predicted Template Modeling
RPA: Replication factor A protein 1
RPS-BLAST: Position-Specific Iterated BLAST
TaSnRK1α: Sucrose non-fermenting-1-related protein Kinase 1α
SD: Synthetic defined
Y2H: Yeast two-hybrid

---

**Supplementary Figure 1.**
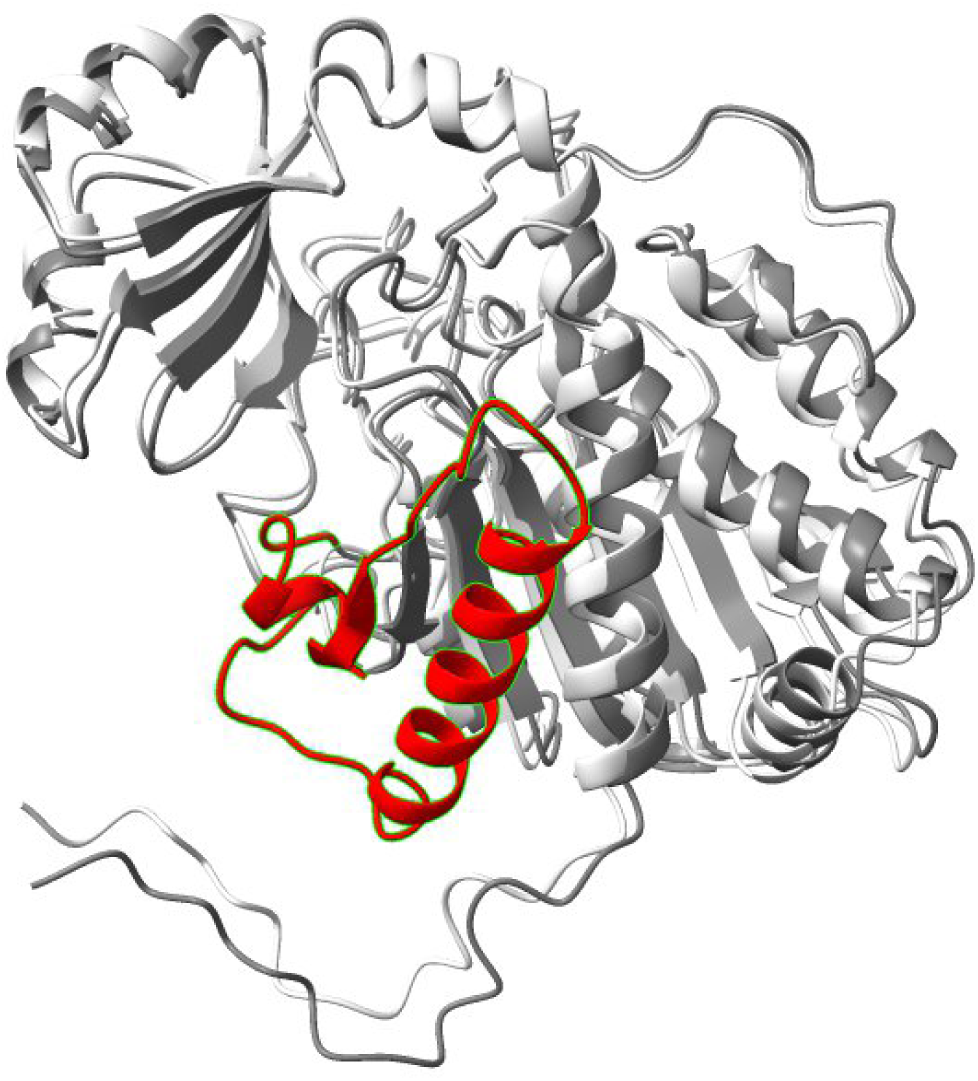
Structural conservation between EFF18 and EFF18_DM. Superimposition of EFF18 (light gray and 45AA used for deletion in red) and EFF18_DM (gray) protein structures with ChimeraX was performed using the *matchmaker* command by creating pairwise sequence alignments and fitting the aligned residue pairs. Root Mean Square Deviation (RMSD) analysis between the conserved or total residues is of 0.372Å and 1.319Å respectively. suggesting that the majority of the structure is well conserved between the two tested proteins.

